# Magnetique: An interactive web application to explore transcriptome signatures of heart failure

**DOI:** 10.1101/2022.07.30.502025

**Authors:** Thiago Britto-Borges, Annekathrin Ludt, Etienne Boileau, Enio Gjerga, Federico Marini, Christoph Dieterich

## Abstract

Despite a recent increase in the number of RNA-seq datasets investigating heart failure (HF), accessibility and usability remain critical issues for medical researchers. We present Magnetique (https://shiny.dieterichlab.org/app/magnetique), an interactive web application to explore the transcriptional signatures of heart failure. We reanalyzed the Myocardial Applied Genomics Network RNA-seq dataset, one of the largest publicly available datasets of left ventricular RNA-seq samples from patients with dilated (DCM) or hypertrophic (HCM) cardiomyopathy, as well as unmatched non-failing hearts from organ donors and patient characteristics that allowed us to model confounding factors. Focusing on the DCM versus HCM contrast, we identified 201 differentially expressed genes and associated pathway signatures. Moreover, we predict underlying signaling networks based on inferred transcription factor activities. To the best of our knowledge, Magnetique is the first online application to provide an interactive view of the HF transcriptome by analyzing differential transcript isoform usage. Finally, another graphical view on statistically predicted RNA-binding protein to target transcript interactions complements the Magnetique web application.

The source code for both the analyses (https://github.com/dieterich-lab/magnetiqueCode2022) and the web application (https://github.com/AnnekathrinSilvia/magnetique) is available to the public. We hope that our application will help users to uncover the molecular basis of heart failure.

## Background

Heart failure (HF) is a complex syndrome, characterized by the interplay of environmental and genetic factors, with a substantial global burden (1). Large-scale gene expression studies have helped to elucidate how transcriptome reprogramming affects underlying gene networks in cardiac remodeling and cardiomyopathies. To this end, the Myocardial Applied Genomics Network (MAGNet) has collected a large quantity of cardiac transcriptomics data, including RNA-sequencing (RNA-seq) and microarray, which have been used to compare transcriptome profiles and gene signatures across a spectrum of HF phenotypes (2)–(3). The MAGNet repository includes left ventricular samples from patients diagnosed with dilated (DCM) and hypertrophic (HCM) cardiomyopathy, as well as from donors with non-failing hearts (NFD), among other etiologies. Most of the MAGNet RNA-seq data are publicly available, and are matched with detailed clinical and demographic information (**Table S01**), which makes it a valuable resource to study the molecular basis of HF. However, these data need to be processed and analyzed, and accessibility, usability, and reproducibility remain a critical issue for life scientists and medical researchers. While repositories such as Gene Expression Omnibus (4) and recount3 (5) provide access to preprocessed data, accessibility to large-scale transcriptomic HF studies with comprehensive patient information remains limited. Poor patient characterization and the lack of standards in experimental and bioinformatics procedures often hinder the extent to which these data can be studied by a broader public.

Here, we provide an interactive web-based solution, called Magnetique (https://shiny.dieterichlab.org/app/magnetique), to explore the transcriptional signatures of HF (**Figure 1**). Magnetique serves as an entry point to 354 MAGNet RNA-seq samples, and provides analyzes with a focus on differential gene expression (DGE) and differential transcript usage (DTU), gene set enrichment analysis (GSEA), protein activity inference, and RNA-binding protein (RBP) interactions to transcript isoforms. Our deployment strategy, combining ShinyProxy and an internal PostgreSQL database, also allows users to repurpose the application with restricted data. In the following, we show how Magnetique can be used to prioritize DTU interactions with regulators, and how ERK1/2 (MAPK3, MAPK1) regulation is central to the differential gene programs observed in DCM versus HCM.

**Figure 1.**
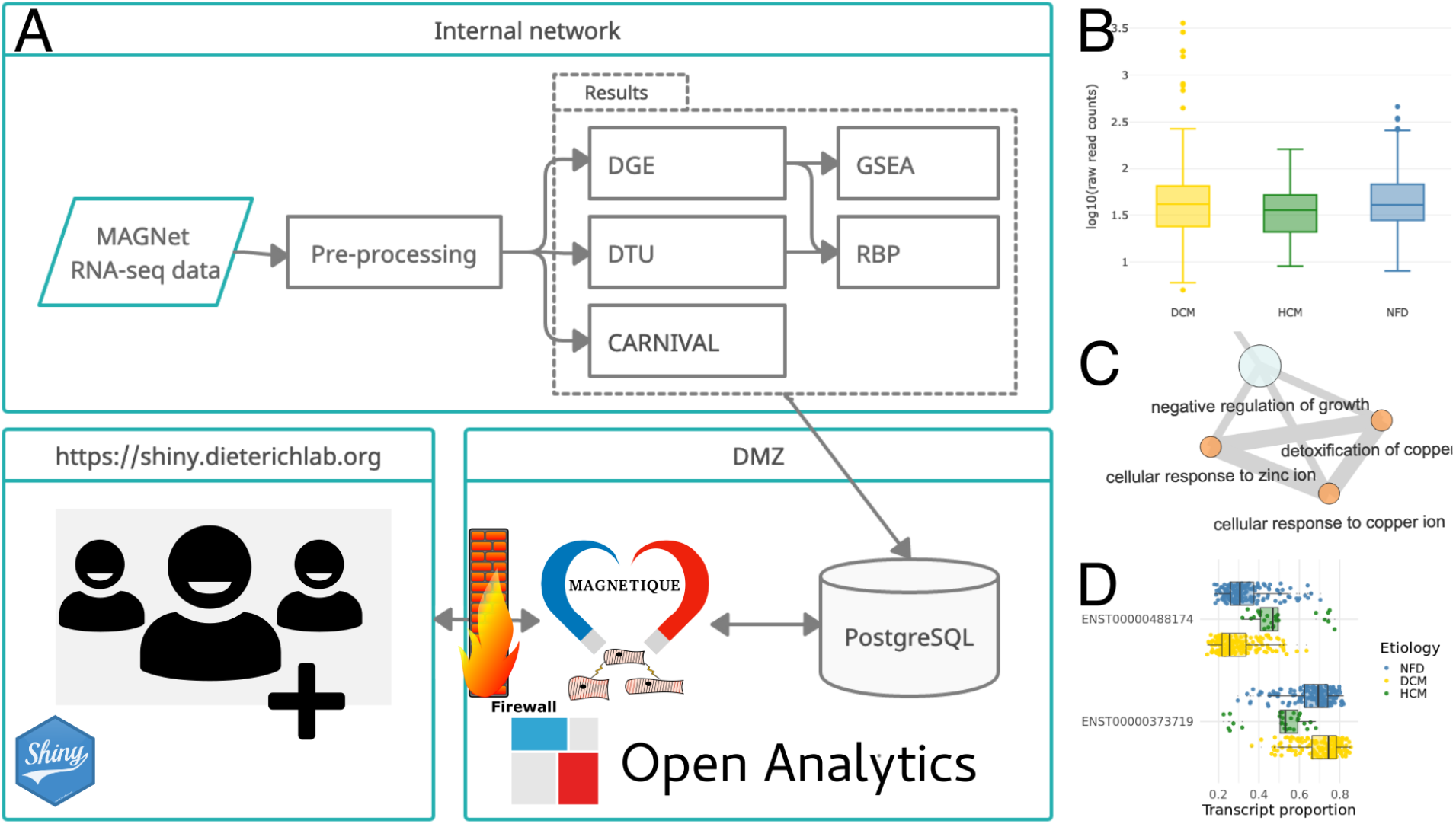
Overview of the Magnetique application. (**A**) RNA-seq data is downloaded, processed, and stored within an internal network. Data is loaded into a PostgreSQL database, which is consumed by the application. The application is served via Open Analytics ShinyProxy on a server located in the demilitarized zone (DMZ) of a high-performance computer cluster located at the Klaus Tschira Institute for Computational Cardiology at Heidelberg University Hospital. The database and ShinyProxy server are protected by a firewall, and external users only have access to the Shiny applications, allowing us to re-use the setup with protected data. Each user instance is served within a Docker container, so user instances are independent and ShinyProxy handles the load on-demand. (**B**). Gene counts for the CXCL3 gene, representing the differential gene expression analysis (DGE), significantly called DGE for the DCMvsHCM contrast. (**C**) Example of an enrichment map for Biological Processes, showing a subgraph of terms originating from a Gene Set Enrichment Analysis (GSEA). (**D**) Differential Transcript Usage (DTU) analysis of the OGT gene locus.

To the best of our knowledge, Magnetique is the first interactive web application to provide DTU analyzes, RBP:RNA interactions, and signaling network inference in the context of HF. It allows researchers with minimum bioinformatics skills to easily browse through the complexity of these data to better understand molecular mechanisms underlying human heart disease.

## Construction and content

### Sequencing data and processing

We used publicly available RNA-seq data from the Myocardial Applied Genomics Network (MAGNet) (GEO: GSE141910), consisting of whole-transcriptomes of left ventricle (LV) tissues from end-stage heart failure (HF) patients due to dilated cardiomyopathy (DCM, n = 165) or hypertrophic cardiomyopathy (HCM, n = 27), and from unmatched non-failing hearts from organ donors (NFD, n = 162).

We first removed adaptors and low-quality bases with Flexbar (v3.5.0) (6). We then identified reads that aligned to human mitochondrial or ribosomal sequences (MT-RNR1, MT-RNR2 and RNA45S5) using Bowtie2 (v2.3.5.1) (7) and discarded them. The remaining reads were aligned to the human genome with STAR 2.6.0c (8). Upon analysis of the read alignment quality control, we observed a high rate of duplicated reads, which we assumed was a technical artifact of the RNA library preparation. We then removed duplicated reads with the MarkDuplicates command from Picard (v2.22.1) followed by samtools (v1.9) (9) to remove reads flagged as PCR or optical duplicates (SAM flag 1024). We generated gene and transcript abundance tables using StringTie2 (v2.1.3b) (10) in the reference guided-mode. For read alignment and transcriptome quantification, we used the ENSEMBL human annotation GRCh38 (v96). **Table S02** shows the versions of the main software packages used for preprocessing, analysis, and application development, as well as links to their repositories.

### Exploratory data analysis, differential gene expression and enrichment analysis

Metadata was downloaded from GSE141910. We also used additional meta data as provided in https://github.com/mpmorley/MAGNet/blob/master/phenoData.csv. Differential gene expression (DGE) analysis was performed using DESeq2 (11) (v1.30.1), accounting for ethnicity (race), age, and sex. We also included two additional latent variables in the model that were found to be correlated with technical artifacts such as DuplicationRate, transcript integrity number (TIN), and library batches (LibraryPool), which found to account for a large proportion of the variance of gene expression (**Figure S01**). Latent variables were determined using surrogate variable analysis (SVA) and confirmed by principal variance component analysis (PVCA) (12) (v1.30.0). We set an adjusted *p*-value ≤ 0.05 as the threshold for calling genes differentially expressed. We produce results for three contrasts: DCM versus HCM, DCM versus NFD, and HCM versus NFD. We used topGO (13) (2.42.0) for gene set enrichment analysis (GSEA) using all expressed genes as background set, nodeSize =10, the “weight01” method, or the wrapper provided by pcaExplorer (14) (v2.16.0), and reported the Fisher’s exact test statistics controlled by the Benjamini & Yekutieli method (15). The mean aggregated log_2_ fold-change scores were obtained with the GeneTonic R package (v1.7.3) (16).

### Differential transcript usage

To perform differential transcript usage (DTU) analysis, we used transcript abundance estimates from StringTie2 transformed to scaledTPM with tximport (v1.18.0) (17). We discarded genes with fewer than 50 read counts or transcripts with usage lower than 0.2 in more than 50 samples, yielding 9,784 genes and 25,730 transcripts (18). The statistical model was similar to the one used for DGE, controlling for ethnicity, sex, age, and the two first components obtained from the surrogate variable analysis. We used DRIMSeq’s implementation of the Dirichlet-multinomial to fit and model the transcript usage (v1.18.0) (19). We reported the likelihood ratio test statistics for the feature level and called significant transcripts with a Benjamini-Hochberg adjusted *p*-value ≤ 0.05. The difference in isoform fraction (DIF) was defined as the difference in usage between two contrasts. The usage was the proportion of a transcript’s expression given the total expression for each gene.

### Prioritization of interactions between transcripts and RNA-binding proteins

We further studied the potential role of RNA-binding proteins (RBPs) on HF by applying Goeman’s Global test on predicted RBP-target transcript pairs using RBP (regulator) expression, target transcription expression, and covariates (20). We used predicted RBP to RNA interactions as described in the oRNAment database (21), yielding 185,052 candidate RBP:RNA interactions for 117 RBPs that were tested for DGE. We only tested RBP encoding genes and RNAs that were tested for DGE and DTU, respectively. It is important to note that the oRNAment database only lists interactions between RBPs and mature RNA sequences and does not capture interactions with introns, which are hardly represented in polyA+ RNA-seq data. To prioritize RBP:RNA interactions, we applied the reverse Global test previously described for prioritization of miRNA to mRNA interactions (22). In short, the *p*-value and direction of change for the association of RBP expression and RNA expression were obtained by iterating over each mRNA and fitting the global test model using the RBP group mean gene expression as a response variable and group mean mRNA expression as data. We selected a FDR corrected *p*-value of 0.05 as the significance cut-off.

### Transcription factor activity-based network inference

We applied CARNIVAL (23) to identify the regulatory processes upstream of TF. CARNIVAL uses Integer Linear Programming to refine protein-protein interaction changes that may drive activation of different gene programs. We collected protein-protein interaction and TF-gene interaction data from OmniPath (24) (v3.3.20) and DoRothEA (v1.7.0) (25), respectively. We filtered OmniPath for genes that were expressed using the edgeR::filterByExpression (v3.3.20) function (26) and for interactions that were signed and directed, yielding a total of 18,098 interactions. For each contrast, we inferred the differential TF activity scores using the VIPER (v1.28.0) method (27) over the DoRothEA regulons and the BiRewire package (v3.26.5) (28) with 1,000 permutations in order to estimate the statistical significance of the enrichment scores. We defined TF with unadjusted *p*-value ≤ 0.1 from the permutation test as differently regulated.

### Web-service deployment

We provided the results of all analyses in a database resource named Magnetique, available at https://shiny.dieterichlab.org/app/magnetique. We used ShinyProxy (29), an open-source solution to deploy our containerized application, and Docker to deliver the web application independently of the operating system. A docker image of the current release for Magnetique is publicly available at https://hub.docker.com/repository/docker/tbrittoborges/magnetique. The application has been tested on Firefox, Chrome, and Safari.

The database backend provides the data to the Shiny application on-demand. The analysis results for the DGE, DTU, Carnival, and RBP analyses, which were stored in SummarizedExperiment objects (30), were processed into tables as serialized JSON versions of the R objects. A PostgreSQL database, described in **Figure S02**, serves the application. This database is protected by a firewall and only accessible from the internal network. Our deployment strategy, combining ShinyProxy and an internal PostgreSQL database, allows users to repurpose the Magnetique application with restricted data, such as patient data, intended for internal use only.

## Utility and discussion

### The Magnetique application

Magnetique is a Shiny application that provides detailed and interactive results for four views: Gene View, Gene set View, Carnival View and RBP:RNA View. We designed Magnetique as a dashboard that is simple to use, targeting a broad audience of biomedical scientists studying the molecular basis of heart failure. Moreover, we included an interactive tour that explains each module and the options available in the application (**Figure S03**). The Gene View (**Figure S04**) comprises the differential DGE analysis and the gene-level summary of the DTU for each contrast. Since the transcriptome profiles were obtained with RNA-seq instead of microarray technology, we could add this novel aspect to our analysis. It includes the tabular result, which links both to the global representation of the DGE and DTU, represented by the volcano plots for both modalities, as well as to detailed statistics for each gene and the involved transcripts. Shifts in transcript usage can imply biological consequences impacting the HF phenotype that are not captured by the DGE analyses.

The Gene set View in Magnetique (**Figure S05**) provides a tabular and graphical network view of overrepresented gene sets, displayed for each of the selected contrasts, to allow a quick identification of the major processes involved, using major GO categories: Biological Process (BP), Molecular Function (MF), and Cellular Component (CC). It includes a heatmap for gene expression signatures in context with the disease etiology and covariates, which allows further stratification of the gene signatures. An enrichment map (31) network details how the different terms relate to one another. Additionally, we show all computed gene expression fold changes, which are stratified by each term.

Using causal reasoning principles, the Carnival View (**Figure S06)** seeks to contextualize large-scale knowledge of protein signaling interactions upstream of significantly differentially regulated TFs, allowing for the study of novel transcriptional regulators in the context of gene expression. Users can bookmark and download their favorite genes and gene sets for later analysis (**Figure S07**). In the RBP:RNA View (**Figure S08**), we graphically present potential RBP:RNA interactions in heart failure, which were called significant when applying the reverse Global test and RNA isoforms that are part of genes tested significantly for DTU (see Construction and Content).

#### Using Magnetique to determine changes in the transcriptome signature of HF

We showcase a comparison of DCM vs HCM patients to identify molecular signatures of different disease etiologies/subtypes in patients with end-stage HF. The DGE results show that out of 41,886 genes tested, 201 were called significantly different (*p*-adjust ≤ 0.05) in the DCMvsHCM contrast (**Figure S09**). 123 differentially expressed genes were common to the three contrasts, while 36 and 42 genes were unique to the DCMvsNFD-DCMvsHCM and HCMvsNFD-DCMvsHCM combinations, respectively (**Figure S10**). These groups of genes are potentially specific markers for the two etiologies, as they distinctly represent the differences between two groups of patients with end-stage cardiomyopathy. Using gene set enrichment analysis to characterize all the differentially expressed genes, we observed Biological Process pathways related to inflammation, specifically neutrophil infiltration, cell-cell signaling and chemokine production, negative regulation of cell growth, and myoblast differentiation. The top 15 Biological Process terms in the context of gene members’ effect sizes are shown in **Figure S11**. Notably, 3 (MSTN, CXCL1 and CXCL3) out of the 10 highlighted genes for the DCMvsHCM contrast are members of the Cytokine-cytokine receptor interaction KEGG pathway (hsa04060). These pathways can be examined in terms of how many members’ genes are shared between pathways using the enrichment map (**Figure S12**).

Differential transcript usage gives a new perspective on genes that are not differentially expressed, but for which changes in transcript usage could lead to phenotypes related to loss-or gain-of-function (32). Interestingly, of the subset of genes that were differentially expressed, only 3 also host transcripts that were DTU. Among the 149 differently used transcripts (**Figure 2**), 107 (79%) were protein-coding transcripts, while 20 (13%) were transcripts with intron retention events. Intron retention is a type of alternative splicing event that occurs when an intron is not spliced out and remains on the mature RNA molecule. This regulatory mechanism has been observed in HF, specifically on genes that are critical to heart function, including MYH7, TNNI3 and TNNT2 (33). Shifts from protein coding isoforms that are translated (messenger RNA) to isoforms without known coding sequence could lead to a net loss-of-function of protein activity. Among the top DTU genes, OGT is a metabolic enzyme that has been linked to cardiac function. The O-linked N-acetylglucosamine (GlcNAc) transferase (OGT) catalyzes a common post-translational modification known for modulating transcriptional regulation (34), heart arrhythmias (35), and heart hypertrophy (36). OGT-202 (ENST00000373719) is the canonical transcript that encodes for the enzyme and is also the primarily used transcript in the NFD and DCM groups. In contrast, OGT-210 (ENST00000488174) is the primary isoform used in HCM condition, and is a transcript with retained intron. Intron retention is a known regulatory process for OGT (37), although this regulation mode for OGT hasn’t yet been implicated in cardiomyopathies.

**Figure 2:**
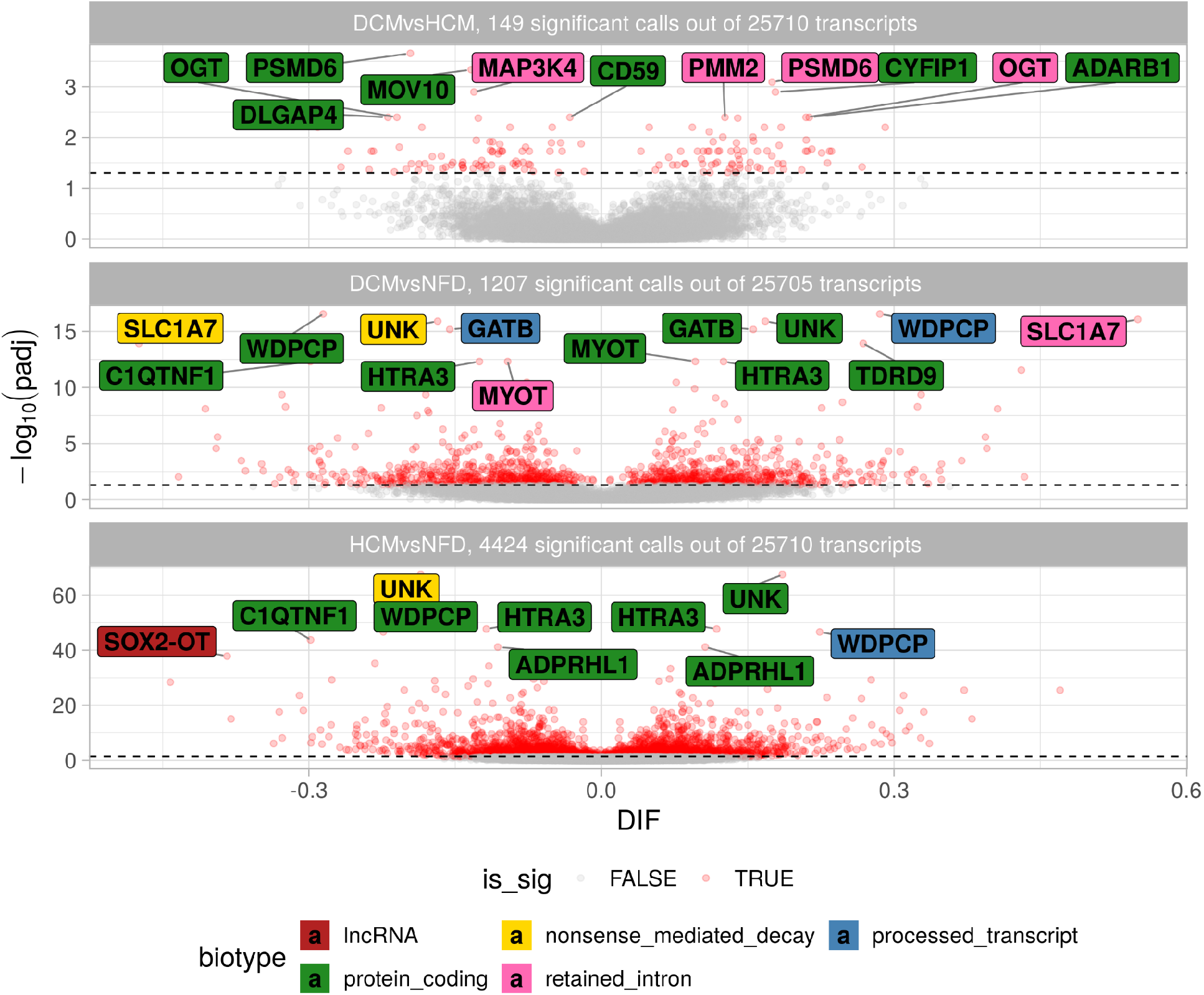
Volcano plot for the DTU analysis. Red circles represent transcripts found significant with an adjusted *p*-value ≤ 0.05. The top ten transcripts for each contrast were named with the gene symbol and colored by the transcript biotype. For the DCMvsNFD and HCMvsNFD contrasts, the volcano shows shifts in isoform usage in several of the top genes, such as OGT, MOV10, ADARB1, GATB, UNK, SLC1A7, HTRA3, ADPRHL1.

In light of the DTU results, we concentrated on post-transcriptional regulation of mRNA by RBP. Interestingly, we found CPEB1 to be the only differentially expressed RBP (**Figure S13**) among 117 tested for DGE and annotated in the oRNAment database (21). CPEB1 regulates mRNA processing and translation by binding to the 3’-UTR of mRNA transcripts, specifically at the cytoplasmic polyadenylation elements. Other members of the CPEB gene family have been linked to heart failure (38). Notably, of the 8,449 CPEB1 targets tested for DTU, 140 were called DTU and 1406 were prioritized by the reverse global test. Six transcript isoforms were both called DTU and have been called significant in the reverse Global test (FDR adjusted *p*-value < 0.05) for CPEB1: ADAMTS9-206 (ENST00000477180), ARHGEF2-202 (ENST00000313695), CPSF4-204 (ENST00000436336), CKAP2-201 (ENST00000258607), METTL23-212 (ENST00000590964), FAM189A2-209 (ENST00000645516), as listed in **Table S4**. Among these, METTL23 is a histone methyltransferase with four isoforms tested for DTU (**Figure S14**) and that has recently been linked to heart congenital disease (39). A difference in the use of an internal exon and the 3’-UTR results in four protein isoforms. CPEB1 binds to METTL23 in its 3’-UTR, which is consistent with the RBP function (40). We observed that CPEB1 gene expression is upregulated in HCM (vs DCM), and the usage of the primary transcript for METTL23, METTL23-212 (ENST00000590964), is drastically reduced in the same experimental group. This finding suggests that further research into the CPEB1:METTL23 interaction, among the other CPEB1:RNA interactions, should be prioritized for the benefit of diagnosis and treatment of cardiomyopathies. Overall, our DTU analysis of the MAGnet RNA-seq data is the first large-scale effort to investigate new markers of HF at the transcript isoform level, which could lead to new hypotheses and interventions (41). These results are accessible through the Magnetique application.

### CARNIVAL network reveals a balance between ERK1 and ERK2 signaling in the failing heart

We applied CARNIVAL to infer the activities of proteins in signaling networks and to dissect potential new regulators involved in HF. Using a signed network of known interactions between proteins, the method propagates the inferred TF activity to upstream regulators for the three comparisons (DCMvsNFD, HCMvsNFD, and DCMvsHCM). **Table S05** lists the number of nodes and edges obtained with CARNIVAL.

ERK1 and ERK2 are extracellular signal-regulated kinases (MAPK3 and MAPK1). Both were central hubs of CARNIVAL networks, with a scaled Kleinberg’s hub centrality score of 1 and 0.356, respectively (**Figure 3**). This centrality score identifies the most influential nodes in a network (42). Interestingly, the signs of the inferred activity for these two kinases have been inferred to be opposite to each other: For the DCMvsHCM contrast, MAPK3 appeared to be consistently less activated in DCM while MAPK1 was more activated in DCM. This suggests a balance between the activities of the two kinases, which can be explained by the differential activity of the PTPRB receptor (receptor-type tyrosine-protein phosphatase beta), which is an upstream regulator for both kinases and was less active in DCM compared to HCM. Both kinases differently regulate a series of TFs and their critical roles in heart homeostasis have been recently reviewed by Gilbert and collaborators (43).

**Figure 3:**
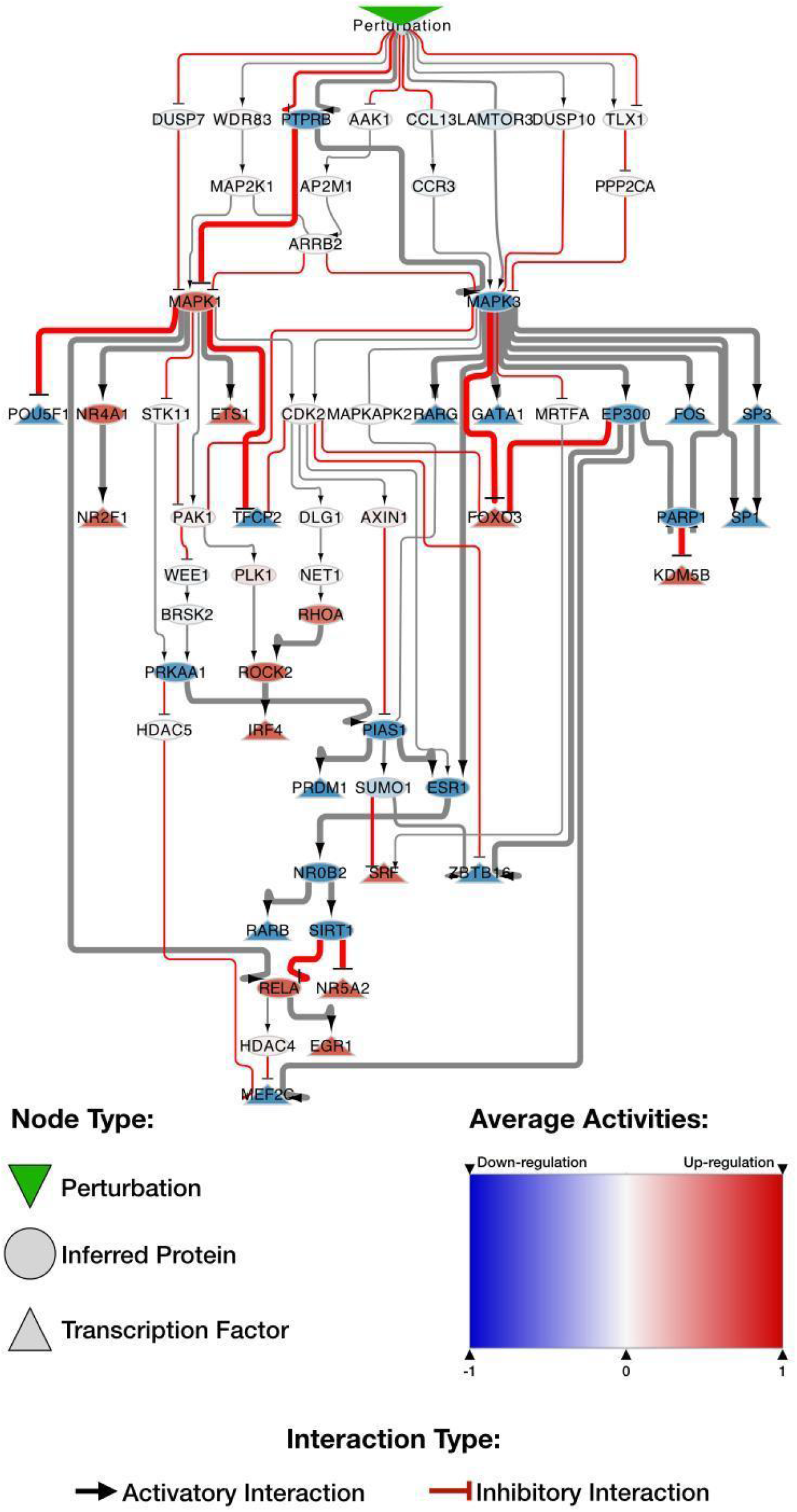
A subset of the CARNIVAL network focusing on the MAPK1 and MAPK3 signaling axes for the DCMvsHCM contrast. Nodes represent members of signaling cascades; round and triangular nodes represent proteins and TFs, respectively. Flat head and arrow head represent interactions that inhibit or activate the downstream ele, respectively. The color of the node represents the inferred protein activity, which ranges from blue (down-regulated) to red (up-regulated). The auxiliary Perturbation node, in the uppermost position, connects all the proteins in the first level of the network. PTPRB, the receptor-type tyrosine-protein phosphatase beta, is likely the primary point for signal transduction. It modulates both MAPK1 (increased activity) and MAPK3 (decreased activity). These protein kinases modulate a series of other factors. Two TFs that are commonly controlled by the MAPK1/MAPK3 signaling axis are MEF2C and EGR1. MEF2C activity is down-regulated by MAPK1 and MAPK3, also known as extracellular signal-regulated kinases (ERKs) 2 and 1, which are central hubs of CARNIVAL networks, with a scaled Kleinberg’s centrality score of 0.356 and 1, respectively. Interestingly, the signs of the inferred activity for these two kinases have been inferred to be opposite to each other: In the DCMvsHCM contrast, MAPK3 appears to be consistently down-regulated while MAPK1 is up-regulated. This suggests a balance between the activities of the two kinases, which can be explained by the differential activity of the PTPRB receptor (receptor-type tyrosine-protein phosphatase beta), which is an upstream regulator for both kinases and is down-regulated. Both kinases differently regulate a series of TFs and their critical roles in heart homeostasis have been recently reviewed by Gilbert and collaborators (via HDAC5) and up-regulated by MAPK3 (via EP300). Conversely, EGR1 activity is controlled by RELA, which is down-regulated by MAPK3 (via SIRT1) and directly up-regulated by MAPK1. In these examples, the activity of TF depends on the balance of the MAPK1/MAPK3 axis.

In fact, the network obtained with CARNIVAL recapitulated part of the known mechanism of action of MAPK3 and MAPK1 in heart failure. For example, ERK1/2 can phosphorylate FOXO3, increasing its nuclear export and thus inhibiting its nuclear effects of activation of pro-apoptotic genes (44). The MAPK family indirectly modulates the activity of the TF MEF2C via the regulation of HDAC4:HDAC5 heterodimer export (45). Conversely, some of the interactions detected with CARNIVAL were novel and thus potentially represent new targets for intervention.

## Discussion and future directions

To the best of our knowledge, there are only a few cardiac-related online databases, such as ANGIOGENES, but their focus is on gene screening, and their data is currently limited to endothelial cells of various tissues (46). The specific challenge of combining multiple gene expression data in the context of HF was recently addressed by the ReHeat portal, also provided as a web-application (47). ReHeat consists of a large meta-analysis of human HF microarray and RNA-seq datasets, referred to as the HF consensus signature. Transcript isoforms or noncoding transcripts are not accessible in ReHeat. In essence, Magnetique is the only publicly interactive application to combine comprehensive RNA-seq analyses into a user-friendly portal. Users with neither technical knowledge nor computational resources can easily access all analyses via four views in graphical and tabular form. Differential transcript usage is often overlooked, despite growing evidence of its role in HF ((48), (41)). Moreover, Magnetique, based on the CARNIVAL inference, allows researchers to identify upstream regulatory signaling pathways that may reveal disease mechanisms. Magnetique thus provides a different approach to analyzing transcriptome signatures in HF. Furthermore, we provide the source code for both the analyses and the web application so that they can be easily replicated and re-used (see Availability of data and materials).

We acknowledge some limitations in the Magnetique application, which will be addressed in future work. For example, Magnetique contains a single dataset, but there are new RNA-seq data sets in the context of HF (e.g., (49)). However, use and access of these data sets within Magnetique need to be typically negotiated with data owners. Second, although the DTU analysis allows the interpretation of shifts in transcript isoform usage, differential splicing methods would have broadened our view of new splicing events, allowing users to identify previously uncharacterized RBP interactions. Finally, we understand that both TFs and RBPs’ activities are also critically regulated at translational and post-translational levels, and plan to consider these levels of regulation, given proteomics and interactome capture data availability, in future versions of Magnetique.

## Conclusions

We re-analyzed a subset of the MAGNet RNA-seq data and provided these results as a web-application that is accessible to a wider audience, addressing issues of data usability and reproducibility of analyses in HF studies. Our analysis focused on resolving technical challenges of the dataset, such as batch effects, and provided a systematic view beyond simple gene expression analyses, supporting hypothesis generation on regulatory processes associated with cardiomyopathies and HF. In summary, Magnetique is the first online application to provide an interactive view of the HF transcriptome at the RNA isoform level and to include transcription factor signaling and RBP: RNA interaction networks. We expect that this may facilitate the uncovering of new diagnostic and therapeutic strategies. Our open-source concept carries over to many other areas of medical research where RNA biology can give new insights beyond gene expression analysis.

## Availability of data and materials

The source code used for the analysis is available at https://github.com/dieterich-lab/magnetiqueCode2022. The source code for the Magnetique web application is available at https://github.com/AnnekathrinSilvia/magnetique. RNA-seq dataset is available through the *Gene Expression Omnibus* with the GSE141910 identifier. A copy for the underlying database is available at https://zenodo.org/record/6854308.

## Authors’ contributions

Conceptualization, C.D.; Methodology, T.B.B., E.B., E.G., and F.M.; Formal Analysis, T.B.B., E.B., and E.G.; Writing - Original Draft, T.B.B., E.B., E.G., and F.M.; Writing - Review & Editing, T.B.B., E.B., E.G., A.L., F.M., and C.D.; Application development, T.B.B., A.L. and F.M.; Funding Acquisition, F.M. and C.D.; Supervision, F.M. and C.D.

## Acknowledgements

We would like to acknowledge Victoria Mauz and Federica Accornero for their feedback on this work. We would also like to thank Harald Wilhelmi for preparing the infrastructure required for Magnetique deployment as well as for technical advice.

