## Supplemental Material for "Magnetique: An interactive web application to explore transcriptome signatures of heart failure"

### Magnetique: A web application for interactive Transcriptome Exploration in Heart Failure

| | Overall, N = 354[1] | DCM, N = 165[1] | HCM, N = 27[1] | NFD, N = 162[1] | p-value[2] |
|---|---|---|---|---|---|
| **Race** | | | | | <0.001 |
| AA | 121 (34%) | 76 (46%) | 1 (3.7%) | 44 (27%) | |
| C | 233 (66%) | 89 (54%) | 26 (96%) | 118 (73%) | |
| **Age** | 56 (48, 62) | 54 (47, 59) | 51 (43, 57) | 57 (51, 65) | <0.001 |
| **Sex** | | | | | 0.021 |
| F | 164 (46%) | 65 (39%) | 11 (41%) | 88 (54%) | |
| M | 190 (54%) | 100 (61%) | 16 (59%) | 74 (46%) | |

| | | | | | |
|---|---|---|---|---|---|
| Weight | 79 (66, 91) | 76 (67, 90) | 72 (60, 83) | 82 (68, 99) | 0.004 |
| Height | 170 (163, 178) | 170 (163, 180) | 170 (160, 180) | 168 (163, 175) | 0.046 |
| Unknown | 1 | 0 | 0 | 1 | |
| HW | 448 (372, 558) | 499 (419, 603) | 448 (366, 559) | 414 (345, 495) | <0.001 |
| Unknown | 7 | 0 | 0 | 7 | |
| LVMass | 247 (199, 322) | 320 (262, 401) | 318 (276, 340) | 214 (179, 278) | <0.001 |
| Unknown | 202 | 119 | 15 | 68 | |
| AFib | 104 (30%) | 67 (41%) | 20 (74%) | 17 (11%) | <0.001 |
| Unknown | 4 | 3 | 0 | 1 | |
| VTVF | 85 (24%) | 70 (43%) | 14 (52%) | 1 (0.6%) | <0.001 |
| Unknown | 2 | 1 | 0 | 1 | |
| Diabetes | 76 (22%) | 39 (24%) | 0 (0%) | 37 (23%) | 0.018 |

| | | | | | |
|---|---|---|---|---|---|
| Unknown | 2 | 0 | 0 | 2 | |
| Hypertension | 85 (24%) | 70 (43%) | 14 (52%) | 1 (0.6%) | <0.001 |
| Unknown | 2 | 1 | 0 | 1 | |
| LVEF | 0.20 (0.15, 0.52) | 0.15 (0.10, 0.20) | 0.25 (0.16, 0.42) | 0.55 (0.50, 0.65) | <0.001 |
| Unknown | 78 | 5 | 1 | 72 | |
| TIN | 70 (61, 73) | 69 (57, 73) | 71 (63, 74) | 71 (65, 74) | 0.019 |
| RIN | 8.50 (8.10, 8.90) | 8.30 (7.80, 8.80) | 8.70 (8.45, 8.85) | 8.70 (8.20, 9.10) | <0.001 |
| Unknown | 6 | 1 | 0 | 5 | |
| DuplicationRate | 0.42 (0.38, 0.52) | 0.42 (0.38, 0.52) | 0.39 (0.35, 0.53) | 0.43 (0.38, 0.51) | 0.6 |
| **SV1** | 0.02 (-0.03, 0.04) | 0.02 (-0.05, 0.04) | 0.04 (-0.02, 0.04) | 0.02 (-0.01, 0.04) | 0.4 |
| **SV2** | -0.02 (-0.04, 0.02) | -0.01 (-0.03, 0.02) | -0.01 (-0.04, 0.02) | -0.02 (-0.04, 0.01) | 0.4 |

[1] Median (IQR); n (%)

| |
|---|
| [2] Kruskal-Wallis rank sum test; Pearson's Chi-squared test |
| Bold highlight features were used in the statistical model. |
| Acronyms: HW: Heart weight; TIN: Transcript Integrity Number: RIN: RNA integrity number; LVMass: Left ventricular mass; AFib: Atrial fibrillation events; VTVF: Ventricular tachycardia/ventricular fibrillation; LVEF: Left ventricular ejection fraction; |

**Table S01**: Patient characteristics and sample metadata.

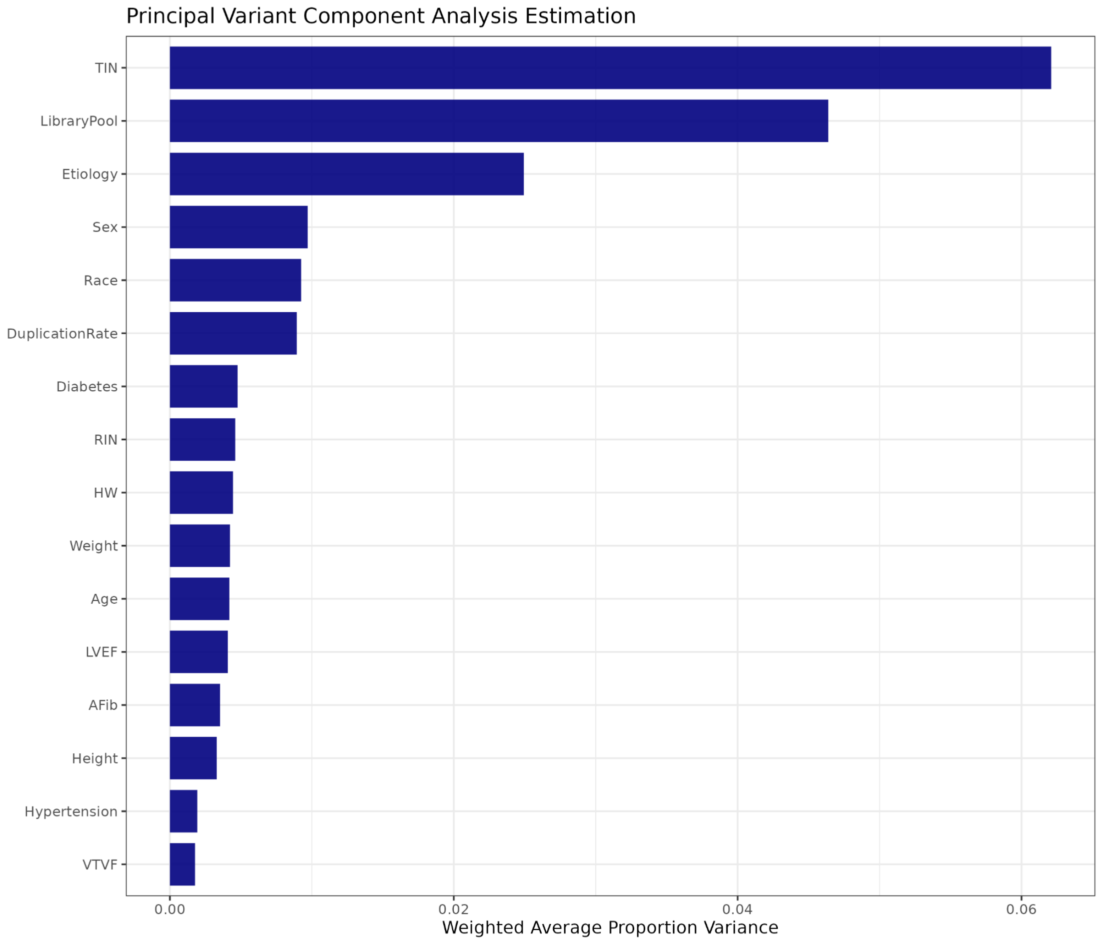


**Figure S01. Principal variant component analysis for covariates.** The majority of the weighted average proportion of variance (0.80) was accounted for by the residuals and it's not shown. IN: Transcript Integrity Number and LibraryPool (batch for library sequence) exhibit greater variation than Etiology.

| **Software** | **Version** | **Repository** |
|---|---|---|
| Flexbar | 3.5.0 | https://github.com/seqan/flexbar |
| Bowtie2 | 2.3.5.1 | https://github.com/BenLangmead/bowtie2 |
| STAR | 2.6.0c | https://github.com/alexdobin/STAR/ |
| Picard tools | 2.22.1 | https://github.com/broadinstitute/picard |
| StringTie2 | 2.1.3b | https://github.com/gpertea/stringtie |

| | | |
|---|---|---|
| sva | 3.38.0 | https://github.com/jtleek/sva |
| pvca | 1.30.0 | https://github.com/dleelab/pvca |
| DESeq2 | 1.30.1 | https://github.com/mikelove/DESeq2 |
| GeneTonic | 1.7.3 | https://github.com/federicomarini/GeneTonic |
| topGO | 2.42.0 | https://bioconductor.org/packages/release/bioc/html/topGO.html |
| Shiny | 1.7.1 | https://github.com/rstudio/shiny |
| tidyverse | 1.3.1 | https://github.com/tidyverse/tidyverse |
| dplyr | 1.0.7 | https://github.com/tidyverse/dplyr |
| ggplot2 | 3.3.5 | https://github.com/tidyverse/ggplot2 |
| GenomicRanges | 1.46.0 | https://github.com/Bioconductor/GenomicRanges |
| rtracklayer | 1.54.0 | https://github.com/lawremi/rtracklayer |
| DRIMSeq | 1.18.0 | https://github.com/gosianow/DRIMSeq |
| DoRothEA | 1.7.0 | https://github.com/saezlab/dorothea |
| edgeR | 3.32.1 | https://bioconductor.org/packages/release/bioc/html/edgeR.html |
| BiRewire | 3.26.5 | http://bioconductor.org/packages/release/bioc/html/BiRewire.html |
| VIPER | 1.28.0 | https://www.bioconductor.org/packages/release/bioc/html/viper.html |
| OmnipathR | 3.3.20 | https://bioconductor.org/packages/release/bioc/html/OmnipathR.html |
| CARNIVAL | 2.4.0 | https://github.com/saezlab/CARNIVAL |
| globaltest | 5.44.0 | https://www.bioconductor.org/packages/release/bioc/html/globaltest.html |

**Table S02. Software used in this work.** The table lists the software, software version, and repository link used on the manuscript.

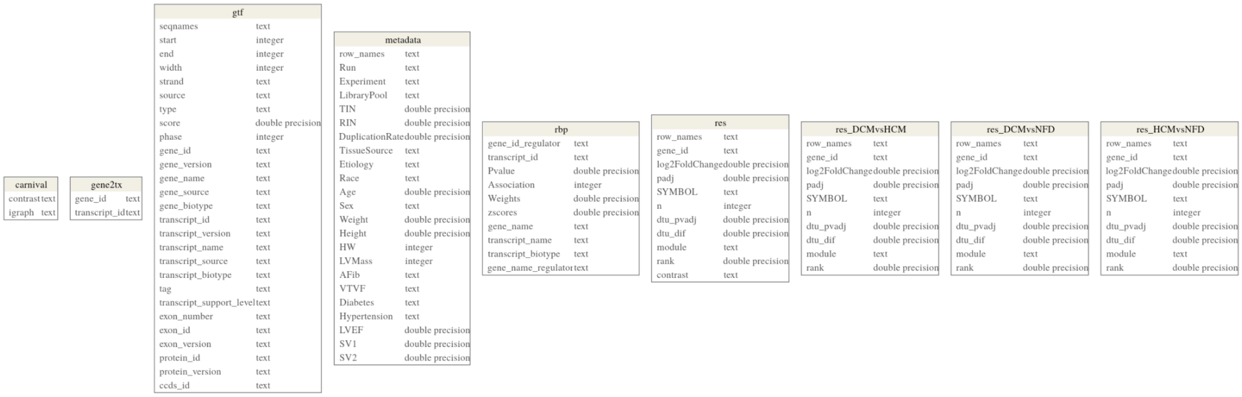


**Figure S02. Magnetique PostgreSQL schema**. The first column in each table represents the column name, and the second column represents the data type. The matrices of features for each patient are represented by three additional tables, named counts (gene counts), dtu fit proportions (fitted transcript proportions), and vst (variance stabilizing transformed gene counts), which were not included in the scheme.

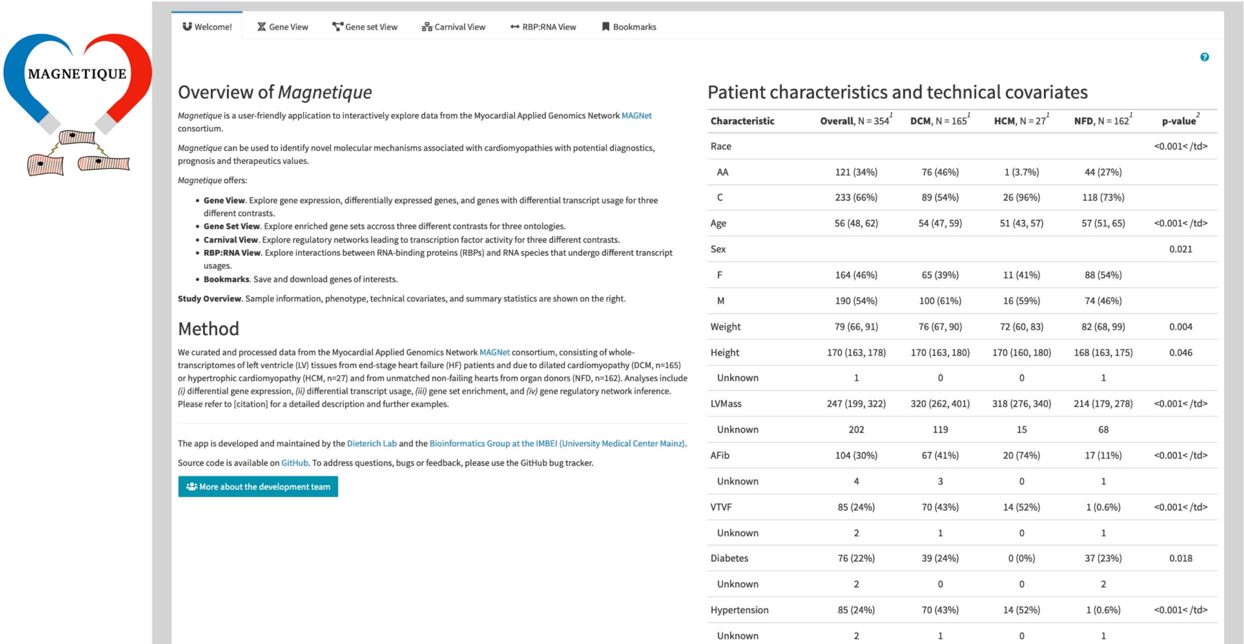


**Figure S03. Partial screenshot of the landing page for the Magnetique application.** The image shows the landing page for the Magnetique application. The different views can be accessed by the multiple tabs at the top. The Welcome tab is followed by the Gene, Gene set, and Carnival views. The help pages are specific to each tab and can be accessed by the interrogation mark at the upper right section on the page (not shown).

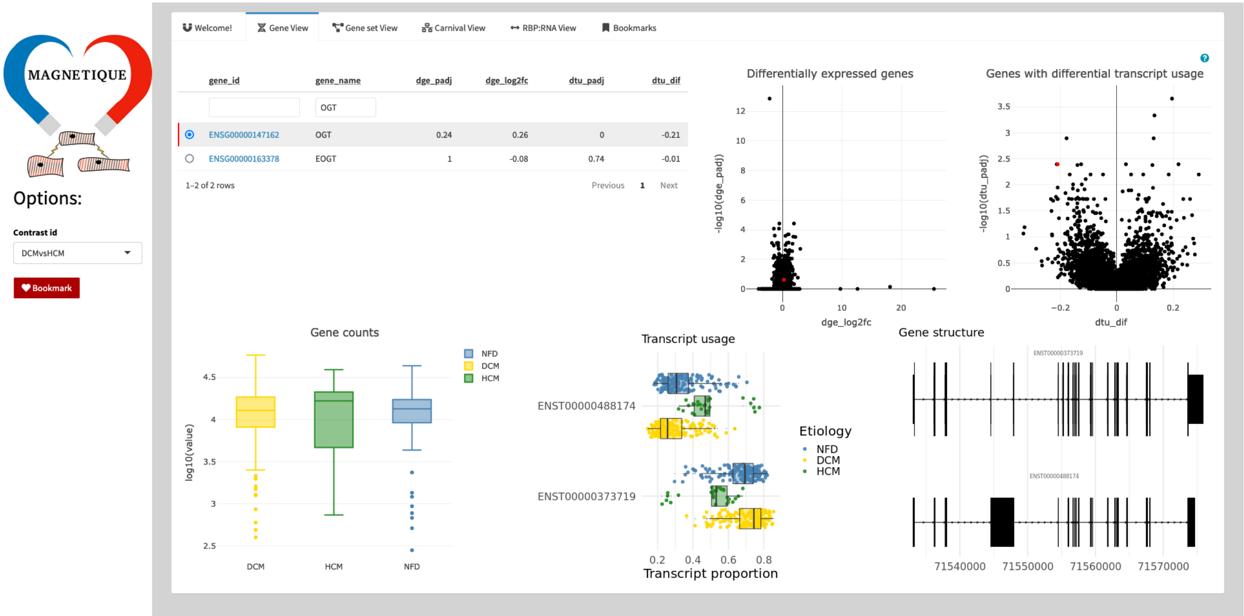


**Figure S04: Gene view page.** This view presents the summaries for the DGE and DTU analyses. In the top left, the table shows the DGE statistics and summary with the minimum adjusted *p*-value and matching DIF for the DTU analysis. In the top center and right, the volcano plots for DGE and DTU, respectively. At the bottom, the summaries for counts and transcript usage and the transcript structure for transcripts that were tested for DTU are triggered once a row in the table is selected. The option in the sidebar (left) allows users to select the contrast.

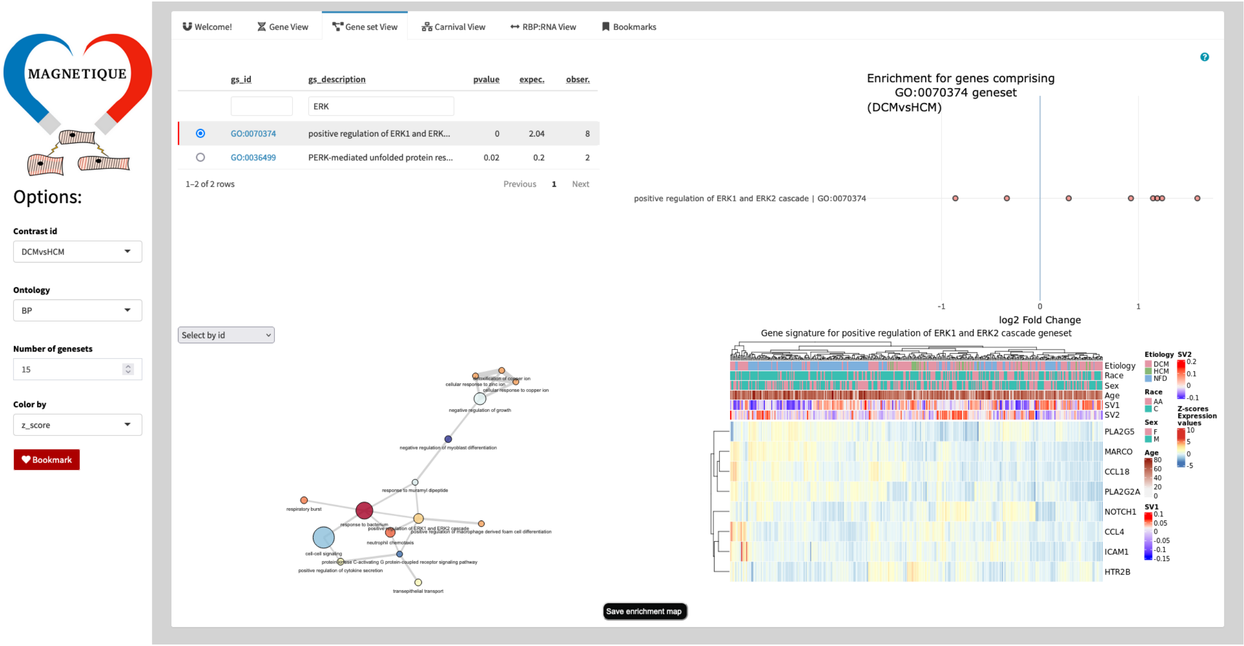


**Figure S05: Gene set view.** The top-left table lists the terms and statistics for the gene set enrichment analysis with TopGo. The top right plot shows the individual log2 fold changes for genes for multiple terms or a single one if a row in the table is selected. The network on the left shows the context among different terms; see Figure S09 for details. The heatmap in the bottom right shows gene signature (z-score transformed gene expression) and covariates used for modeling for a given term, once selected from the table. Multiple options can be set on the side panel.

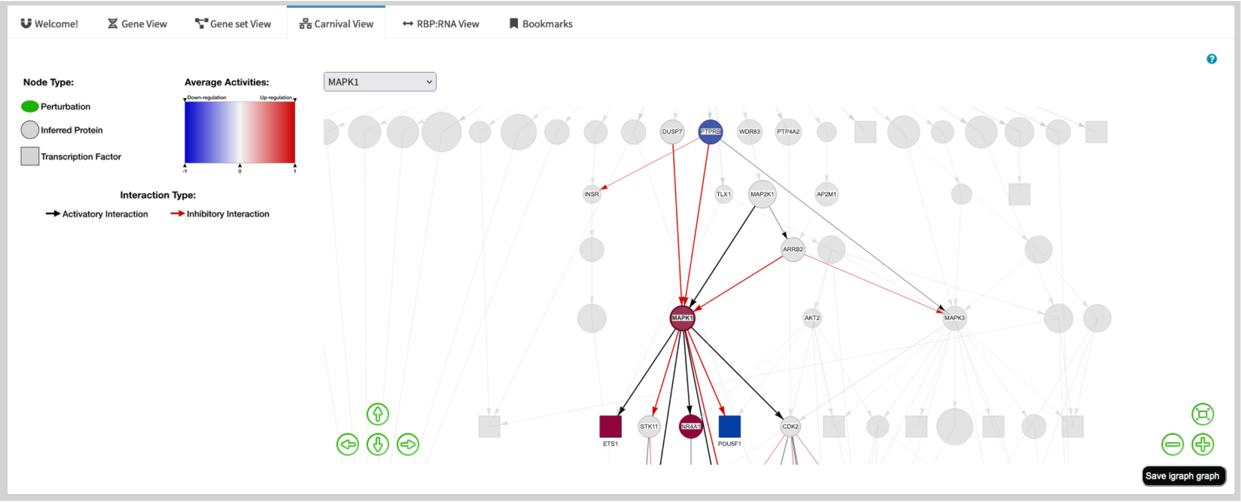


**Figure S06: Carnival View.** The network shows multiple levels of protein signaling networks and inferred protein activities given the gene expression signature of a contrast (selected in the options). Regulatory interactions represented by the edges (edges) can either activate or inhibit targets. Transcription factors are represented by rectangular nodes.

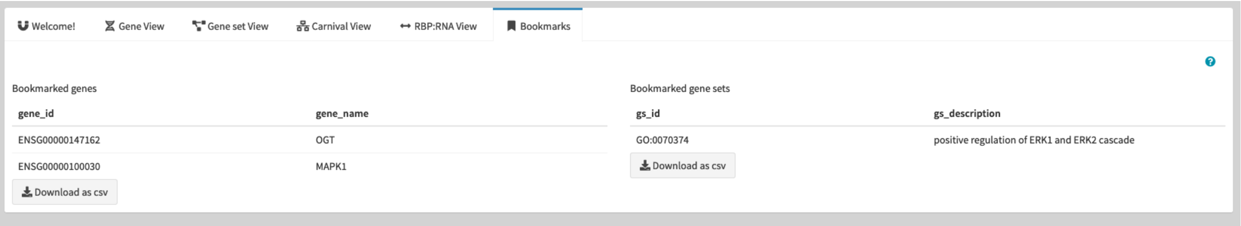


**Figure S07: Bookmarks tab.** The bookmarks tab shows the bookmarked genes from the Gene View or Carnival View and gene sets from the Gene set View that were selected with the bookmark button. The lists can be downloaded with the "*download as csv"* button.

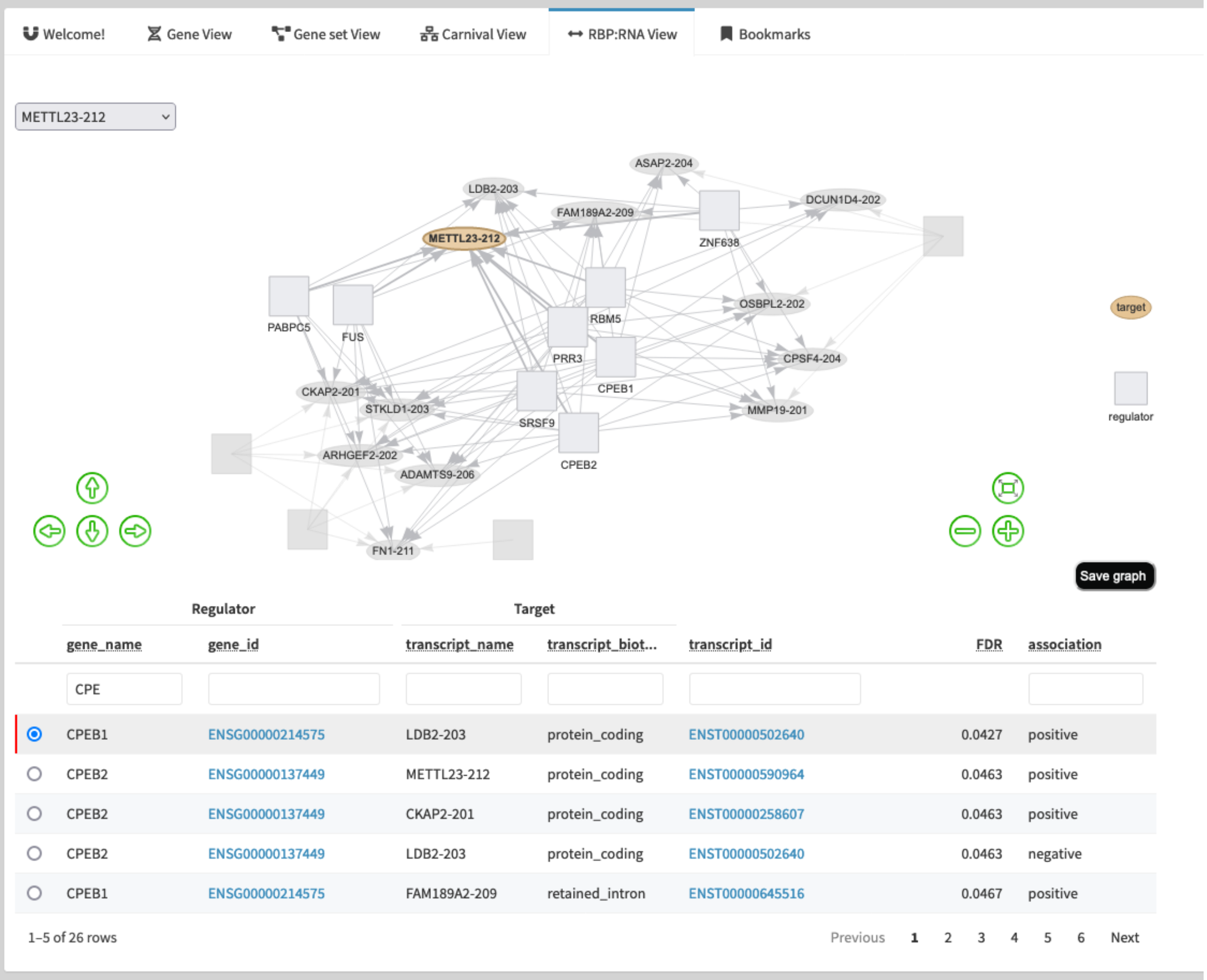


**Figure S08 Overview of RBP:RNA View.** This view lists the interactions between RBP (regulator) and transcripts isoforms (targets) for genes that host significant DTU events. The view filters for CPEB1.

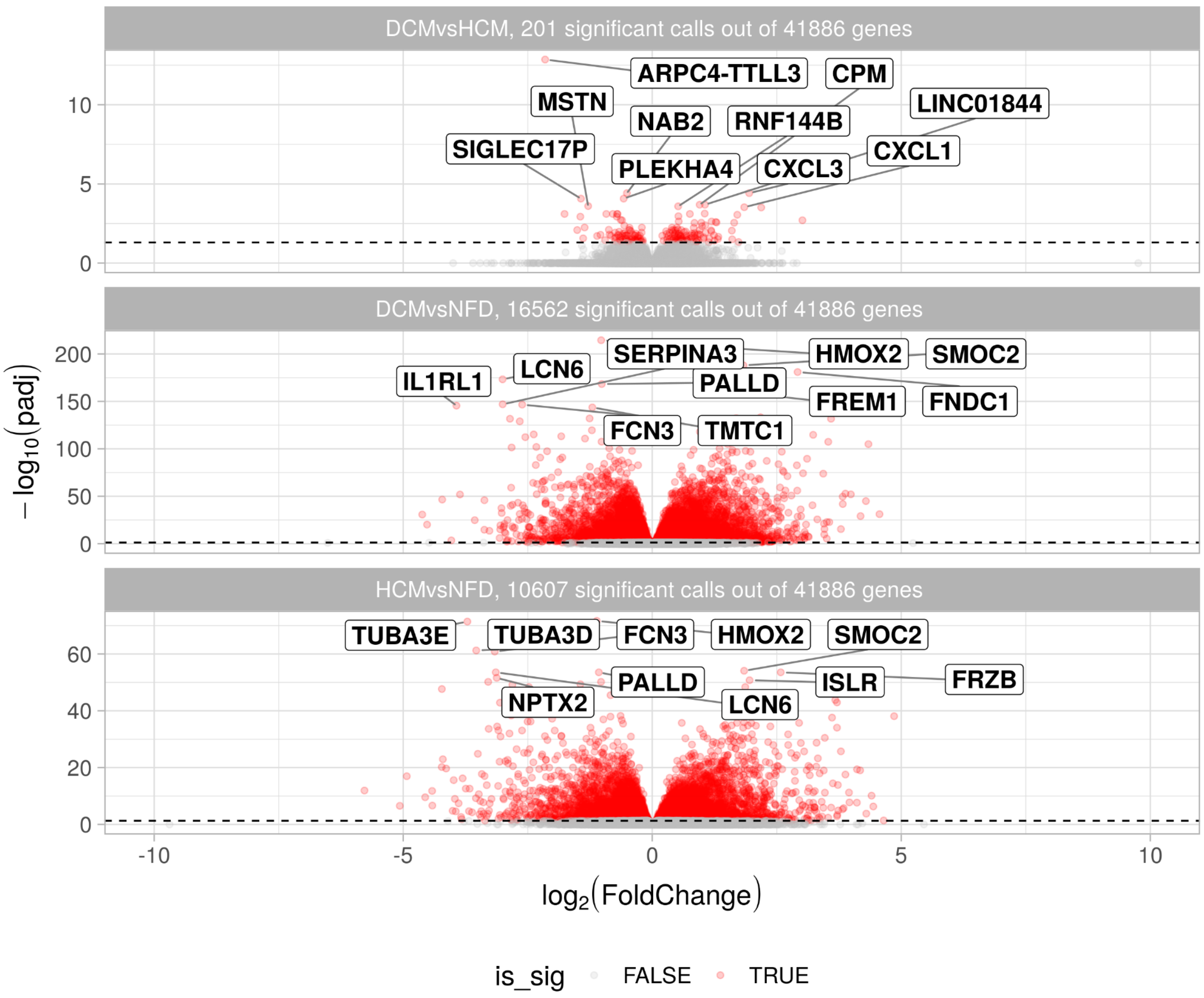


**Figure S09: Volcano plot for the DGE analysis.** Red circles represent genes called significant (adjusted p-value ≤ 0.05). Top ten genes for each contrast were named with the gene symbol.

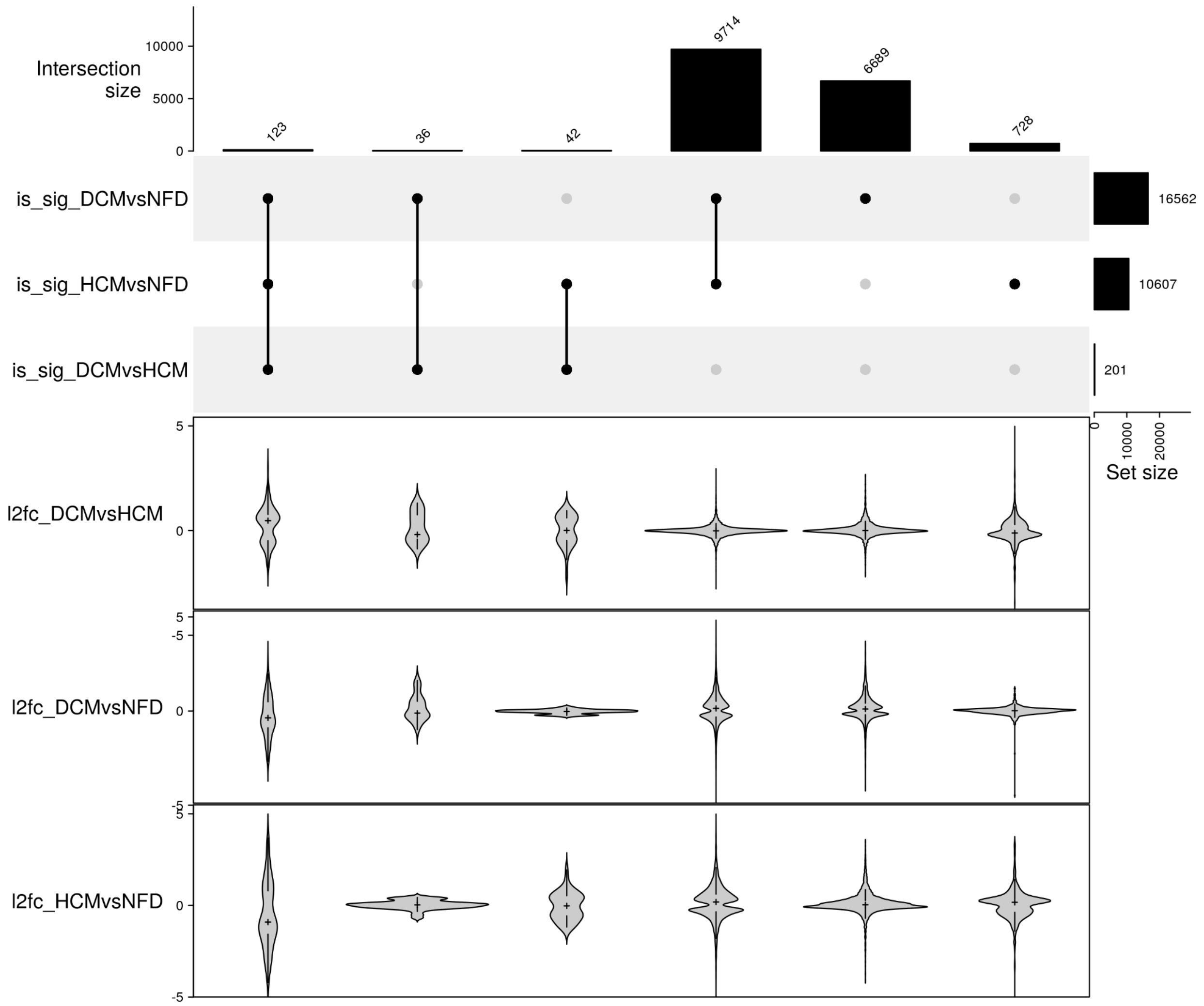


**Figure S10: Combinations of differentially expressed genes for among contrasts.** A total of 123 genes are commonly differentially expressed by the three contrasts. Genes in this combination have a median $\log_2$(FoldChange) of 0.48, and so are upregulated.

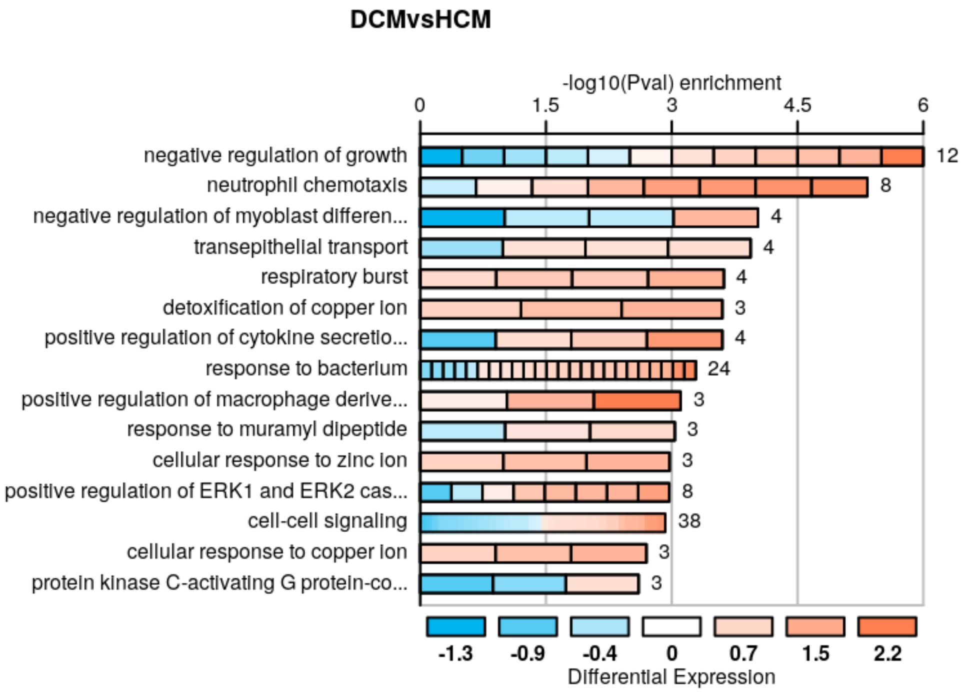


**Figure S11**: **The top 15 Biological Process terms ranked *p*-value for the DCMvsHCM comparison.** Each cell represents a gene and the cell colors show the effect sizes for the DGE analysis, as annotated in the bottom of the image.

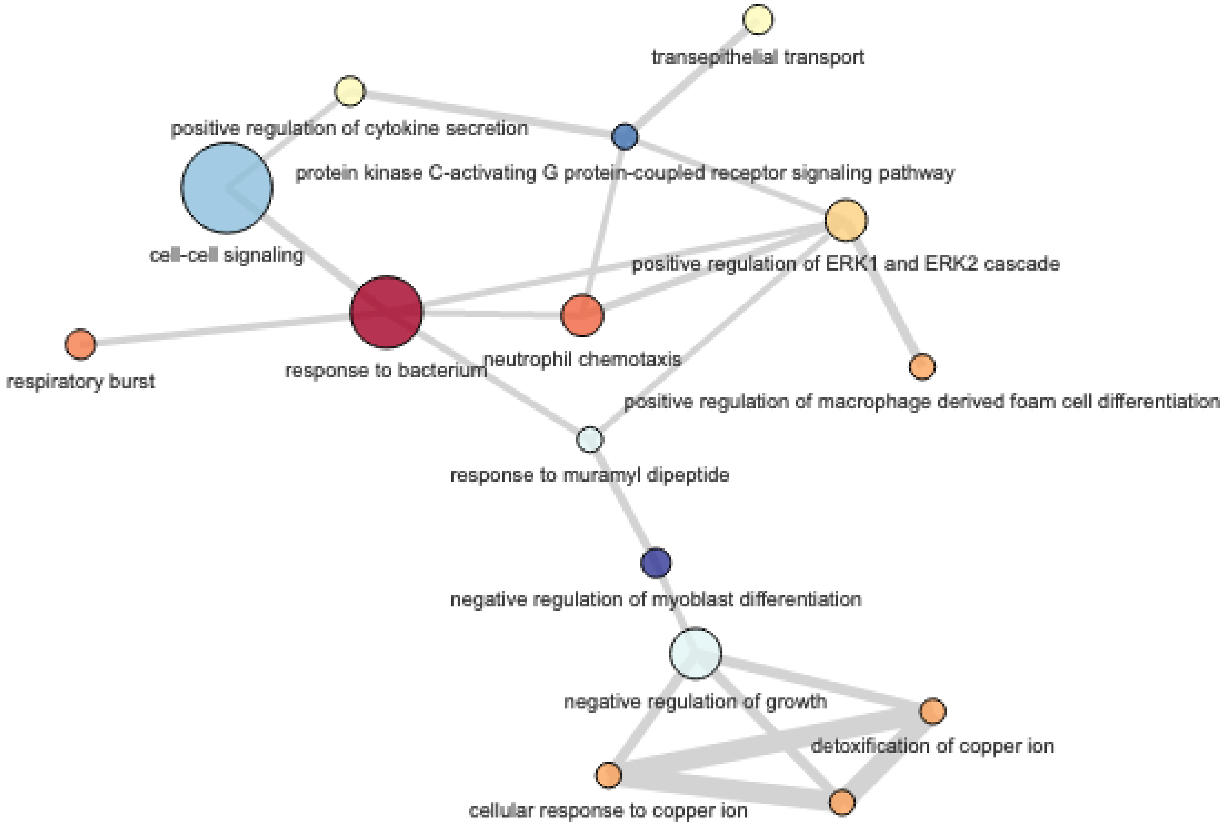


**Figure S12: The enrichment map for top biological process terms called significant in the GSEA.** Node sizes represent the number of genes per term, and node color represents the mean DGE effect sizes (z-score transformed) for gene members of each gene set; red nodes have more genes highly expressed in DCM, while blue nodes have more genes highly expressed in HCM. Edge thickness represents the amount of overlapping gene members within the gene sets. The cell-cell signaling pathway was the term with the highest number of DGE genes. Negative regulation of myoblast differentiation (GO:0045662, more genes highly expressed in HCM) and response to bacterium (GO:0009617, more genes highly expressed in DCM) are, respectively, terms for each with the summarized (mean z-score transformed) gene expression effect size are more extreme.

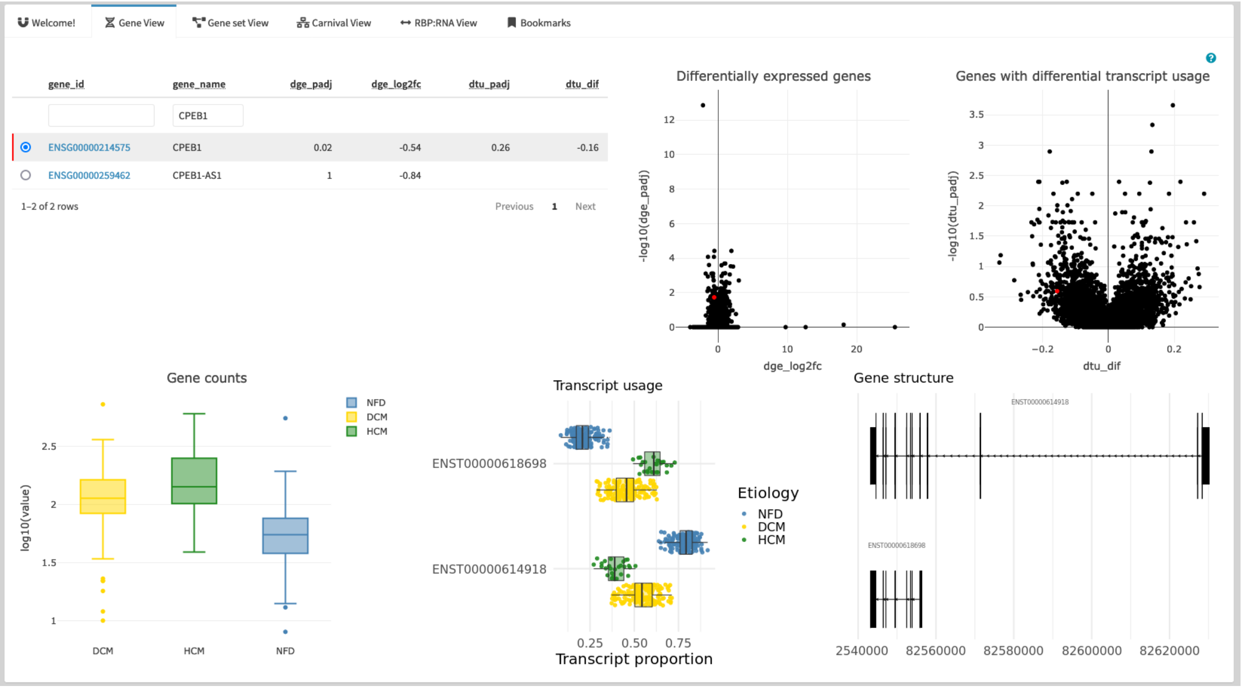


**Figure S13 Overview of DGE and DTU for CPEB1.** CPEB1, the Cytoplasmic Polyadenylation Element Binding Protein 1. It is differentially expressed for DCMvsHCM comparison, but also undergoes different transcript usage for DCMvsNFD and HCMvsNFD. The two disease condition show the same pattern for isoform usage: the primary use of CPEB1-211 (ENST00000615198) while the longer isoform (CPEB1-209 - ENST00000614918) is preferably used in the NFD.

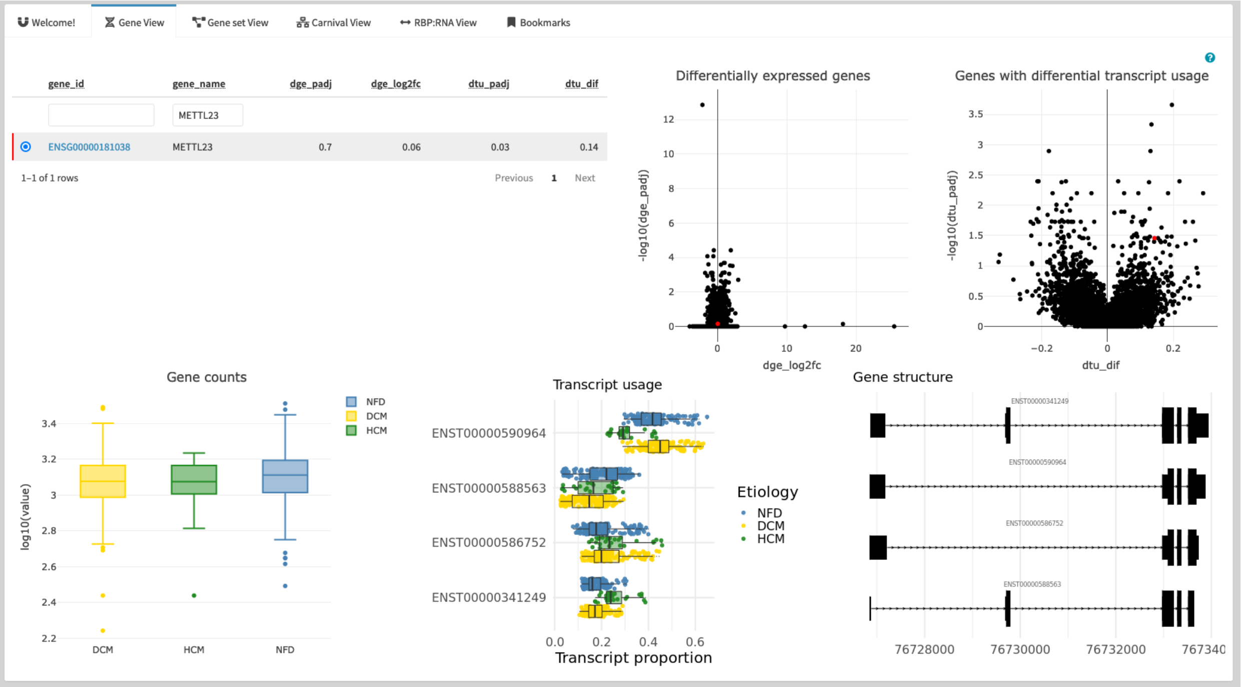

**Figure S14 Overview of DGE and DTU for METTL23.** Histone-arginine methyltransferase METTL23 is one of the targets that was prioritized by reverse global test analysis and also undergoes DTU.

| Gene_id regulator | Gene_name regulator | Transcript_name Target | Transcript_id Target | Transcript_biotype Target | FDR | Association |
|---|---|---|---|---|---|---|
| ENSG00000214575 | CPEB1 | FAM189A2-209 | ENST00000645516 | retained_intron | $4.7 \times 10^{-2}$ | + |
| ENSG00000214575 | CPEB1 | METTL23-212 | ENST00000590964 | protein_coding | $4.7 \times 10^{-2}$ | - |
| ENSG00000214575 | CPEB1 | CKAP2-201 | ENST00000258607 | protein_coding | $4.7 \times 10^{-2}$ | - |
| ENSG00000214575 | CPEB1 | CPSF4-204 | ENST00000436336 | protein_coding | $4.7 \times 10^{-2}$ | - |
| ENSG00000214575 | CPEB1 | ARHGEF2-202 | ENST00000313695 | protein_coding | $4.9 \times 10^{-2}$ | + |
| ENSG00000214575 | CPEB1 | ADAMTS9-206 | ENST00000477180 | retained_intron | $4.9 \times 10^{-2}$ | - |

**Table S04: Targets of CPEB1 with FDR ≤ 0.05 for the reverse global test and adjusted *p*-value < 0.05 for DTU.**

| | DCMvsHCM | DCMvsNFD | HCMvsNFD |
|---|---|---|---|
| Nodes (proteins) | 111 | 142 | 92 |
| Edges (interactions) | 201 | 265 | 164 |

**Table S05: Number of components for the network solutions obtained by CARNIVAL.**
